# Ocean-scale variation in migration schedules of a long-distance migratory seabird is fully compensated upon return to the breeding site

**DOI:** 10.1101/2023.05.27.542544

**Authors:** Rob SA van Bemmelen, Børge Moe, Hans Schekkerman, Sveinn Are Hansen, Katherine RS Snell, Elizabeth M Humphreys, Elina Mäntylä, Gunnar Thor Hallgrimsson, Olivier Gilg, Dorothée Ehrich, John Calladine, Sjúrður Hammer, Sarah Harris, Johannes Lang, Sölvi Rúnar Vignisson, Yann Kolbeinsson, Kimmo Nuotio, Matti Sillanpää, Benoît Sittler, Aleksandr Sokolov, Raymond H.G. Klaassen, Richard A. Phillips, Ingrid Tulp

**Affiliations:** Wageningen Marine Research, Haringkade 1, 1976 CP IJmuiden, the Netherlands. Current address: Waardenburg Ecology, Culemborg, the Netherlands and Wageningen Marine Research, IJmuiden, the Netherlands; Norwegian Institute for Nature Research (NINA), Trondheim, Norway; SOVON Vogelonderzoek Nederland, Nijmegen, the Netherlands; Norwegian Institute for Nature Research (NINA), Tromsø, Norway; University of Copenhagen, Copenhagen, Denmark & Max Planck Institute of Animal Behavior, Radolfzell, Germany; British Trust for Ornithology (BTO), Scotland, Stirling University Innovation Park, Stirling, FK9 4NF, United Kingdom; Applied Zoology/Animal Ecology, Institute of Biology, Freie Universität Berlin, Berlin Germany; Institute of Entomology, Biology Centre of the Czech Academy of Sciences, České Budějovice, Czech Republic; Faculty of Sciences, University of South Bohemia, České Budějovice, Czech Republic & Section of Ecology, Department of Biology, University of Turku, Turku, Finland; Department of Life and Environmental Sciences, University of Iceland, Reykjavik, Iceland; 1UMR 6249 Chrono-Environnement, CNRS, Université de Bourgogne Franche Comté, 25000 Besançon, France, 2Groupe de Recherche en Ecologie Arctique, 16 rue de Vernot, 21440 Francheville, France; UiT The Arctic University of Norway, Tromsø, Norway; British Trust for Ornithology (BTO), Scotland, Stirling University Innovation Park. Stirling, FK9 4NF, United Kingdom; 1Faroese Environment Agency, Traðagøta 38, Argir, FO-165, Faroe Islands, Faroe Islands; 2University of the Faroe Islands, Faculty of Science and Technology, Vestarabryggja 15, FO-100 Tórshavn, Faroe Islands; British Trust for Ornithology (BTO), The Nunnery, Thetford, Norfolk IP24 2PU, United Kingdom; 1University of Giessen, Giessen, Germany; 2Groupe de Recherche en Ecologie Arctique, 16 rue de Vernot, 21440 Francheville, France; Sudurnes Science and Learning Center, Suðurnesjabær, Iceland; Northeast Iceland Nature Research Centre, Husavik, Iceland; Pori Ornithological Society, Pori, Finland; Environmental Agency, Pori, Finland; Pori Ornithological Society, Pori, Finland; 1University of Freiburg, Freiburg, Germany; 2Groupe de Recherche en Ecologie Arctique, 16 rue de Vernot, 21440 Francheville, France; Arctic research station of Institute of plant and animal ecology, Ural branch, Russian Academy of Sciences, Labytnangi, Russia; Conservation Ecology Group, Groningen Institute for Evolutionary Life Sciences (GELIFES), Groningen University, Groningen, Netherlands; British Antarctic Survey (BAS), Natural Environment Research Council (NERC), Cambridge, United Kingdom; Wageningen Marine Research, IJmuiden, the Netherlands

**Keywords:** Arctic Skua, Parasitic Jaeger, *Stercorarius parasiticus*, migratory connectivity, phenology, annual cycle, carry-over effects

## Abstract

**Background:** Migratory birds generally have tightly scheduled annual cycles, in which delays can have carry-over effects on the timing of later events, ultimately impacting reproductive output. Whether temporal carry-over effects are more pronounced among migrations over larger distances, with tighter schedules, is a largely unexplored question.

**Methods:** We tracked individual Arctic Skuas *Stercorarius parasiticus*, a long-distance migratory seabird, from eight breeding populations between Greenland and Siberia using light-level geolocators. We tested whether migration schedules among breeding populations differ as a function of their use of seven widely divergent wintering areas across the Atlantic Ocean, Mediterranean Sea and Indian Ocean.

**Results:** Breeding at higher latitudes led not only to later reproduction and migration, but also faster spring migration and shorter time between return to the breeding area and clutch initiation. Wintering area was consistent within individuals among years; and more distant areas were associated with more time spent on migration and less time in the wintering areas. Skuas adjusted the period spent in the wintering area, regardless of migration distance, which buffered the variation in timing of autumn migration. Choice of wintering area had only minor effects on timing of return at the breeding area and timing of breeding and these effects were not consistent between breeding populations.

**Conclusion:** The lack of a consistent effect of wintering area on timing of return between breeding areas indicates that individuals synchronize their arrival with others in their population despite extensive individual differences in migration strategies.

## Introduction

Stages in the annual cycle of animals are intricately linked and the timing of each event can have cascading effects on the timing of subsequent events, usually referred to as carry-over effects [1,2]. Temporal carry-over effects may be particularly pronounced in animals that undertake long and challenging migrations between their breeding and wintering areas, where delays in one phase of the annual cycle can lead to a temporal mismatch with resource availability in subsequent phases, ultimately affecting survival and reproductive success [3,4]. The degree to which delays can be buffered, preventing amplification of carry-over effects across time, may vary as a function of the tightness of scheduling within the annual cycle [5]. Quantifying the geographical variation in timing of annual cycles and carry-over effects is therefore important for our understanding of how phenology drives variation in life-history traits [6]. In addition, considering that daily survival rates may differ between stages of the annual cycle [7,8], variation in annual schedules can drive meta-population dynamics [2,9,10]. Insight in the drivers of meta-population dynamics is important to inform location-specific conservation efforts.

Population-level differences in the scheduling of the annual cycle may arise as a function of the location of the breeding and wintering areas [11,12] and the conditions experienced there [13]. Annual schedules are generally shifted to later in the season at higher breeding latitudes, where the optimal time for reproduction is associated with a later time of return to the breeding site [18]. In turn, this affects the duration and timing of subsequent phases of the annual cycle, such as migration, given that the degree of seasonality in resources and photoperiod leads to differences in optimal migration speed [19,20]; however, this is little studied. In addition to an effect of the breeding area, the timing of annual cycles may also depend on the choice of wintering area and migration routes, as more distant non-breeding areas require longer travel time. As such, unless departure is earlier or travel speeds higher, arrival will be later [21,22]. When fueling conditions in the wintering area itself or along the migration route are sub-optimal, delays can arise as a result of choice of wintering area. These delays can subsequently carry-over to timing of return to the breeding area and egg laying [23,24] and can reduce survival [25]. Thus, temporal carry-over effects are expected to be more pronounced when distances between breeding and non-breeding areas are greater, and there is less leeway in the annual cycle to compensate for delays.

We define the *net* carry-over effect as the timing of an event for an individual or group relative to the population mean for a sequence of events (figure 1a). The *strength* of temporal carry-over effects can be conceptualized as the degree to which timing of an event affects the scheduling of a following event (figure 1a-c). If there is full compensation for a delay, each day lost in one phase translates to one less day spent in the next phase, resulting in a similar timing of the following event. In the case of complete carry-over of delays, the timing of the next event is delayed by a day, i.e. there is no compensation. Delays may also be amplified at later stages, i.e. each day of delay translates to more than a day delay in following events.

**Figure 1:**
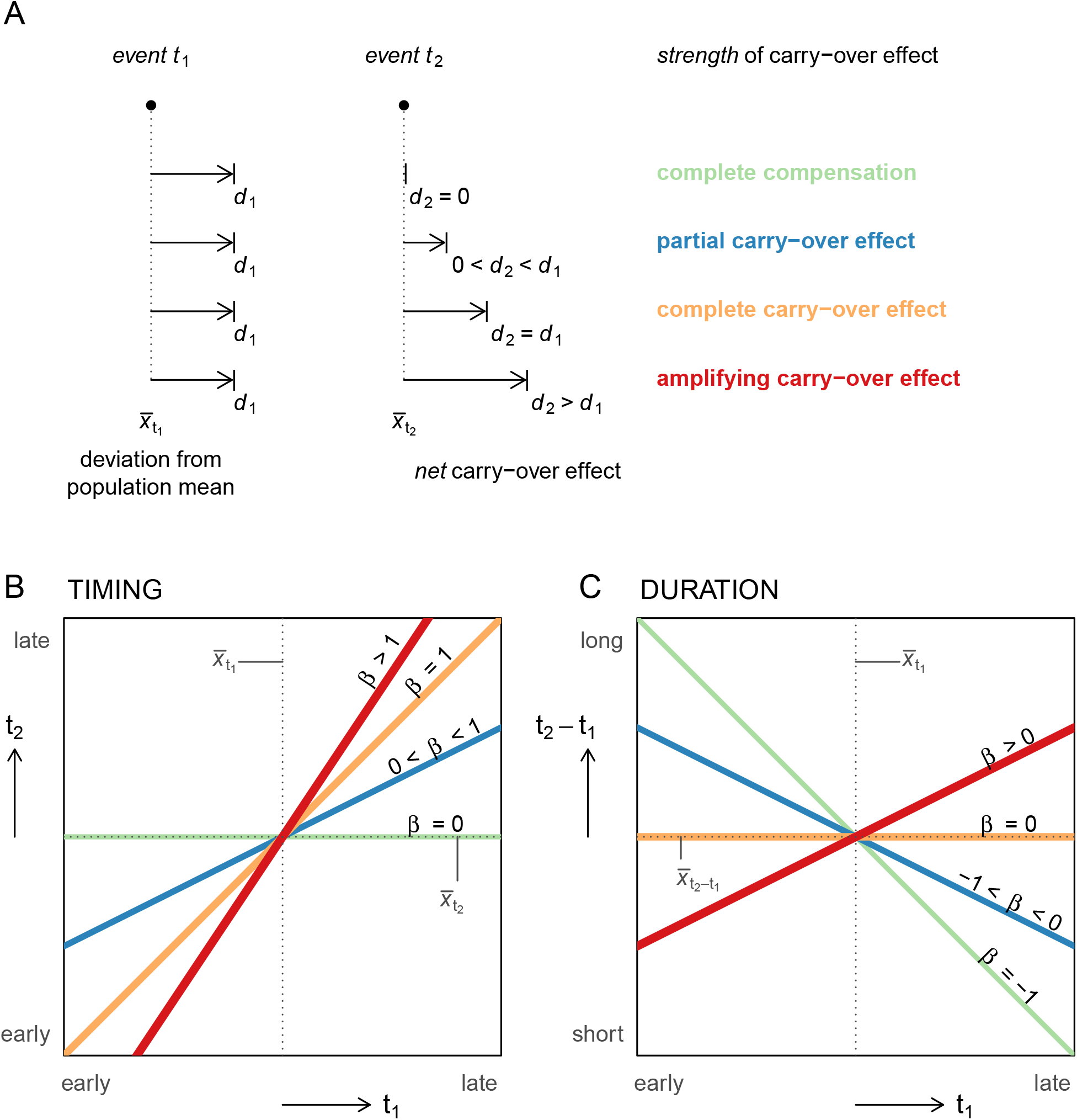
Conceptual framework of the two interrelated measures of carry-over effects between events in the annual cycle. The framework assumes the population mean represents the overall optimal timing, while throughout the range of timings different advantages and disadvantages can be attributed. (A) Here an example delay (*d*_1_) relative to the population mean timing (*x̄*_*t*_1__) of event 1 affects the timing of a subsequent event *t*_2_ relative to the population mean *x̄*_*t*_2__ to different degrees. The resulting degree of advancement or delay in event (*t*_2_) is the *net* of carry-over effects and is described by its *strength* as partial, complete, or amplified carry-over effect, or a complete compensation. The *strength* of the carry-over effect can be quantified as the slope between *t*_1_ and *t*_2_ (B), or, between *t*_1_ and the duration of the period between *t*_1_ and *t*_2_ (C).

Although long-distance migration has evolved in many taxonomic groups [26], geographical variation in the timing of annual cycles and carry-over effects have mainly been studied in single breeding populations of migratory birds [13,27,28], that use a single wintering area or multiple wintering areas at similar latitudes [11,12,15] or in multiple breeding populations that breed at similar latitudes [29]. Ideally, quantifying and testing the relative effects of breeding and wintering location on the timing of annual schedules and on carry-over effects requires substantial geographic variation in both breeding and wintering areas and mixing of individuals from different breeding populations within wintering areas and *vice versa* (weak migratory connectivity [30]). Such studies are rare, because of the challenges of performing large-scale tracking studies and because few species show sufficient variation in both breeding and wintering latitudes. Hence, the extent to which the timing of events and carry-over effects in the annual cycle of migratory birds is driven by both breeding and wintering locations is largely unresolved [31].

We studied carry-over effects on the timing of different phases in the annual cycle of a long-distance migratory seabird, the Arctic Skua *Stercorarius parasiticus*, from multiple breeding populations. Its circumpolar breeding distribution ranges from northern temperate to high arctic zones [[32]; figure 2]. Based on ring recoveries and field sightings, the main wintering areas in the Atlantic are thought to span productive continental shelf and pelagic areas across a large (latitudinal) range, from Iberia to Patagonia and South Africa [32,33]. By tracking the full annual cycle of many individuals over several years from multiple breeding sites ranging from East Greenland to West Siberia (59°N to 79°N in latitude and 24°W to 69°E in longitude), we present the full extent of their wintering distribution. We quantify the migratory connectivity [30] and the consistency of individuals from year to year in their migration strategy. We quantify the relative effects of breeding and wintering areas on the timing of key events and duration of intervening periods in the annual cycle. Based on the northern hemisphere (breeding area) viewpoint on annual schedules, the non-breeding area is referred to as wintering area throughout. Using the variation in annual schedules, we study compensation versus carry-over of variation in timing between subsequent events, and how these depend on breeding and wintering areas. Later timing of events and tighter schedules are predicted to be associated with more northerly breeding areas [15,16]; later timing is also expected to lead to different allocation of time to phases of the annual cycle. Time spent in more distant wintering areas is expected to be shorter due to longer time spent on migration. Among different annual schedules, we expect carry-over effects to arise particularly among schedules with less leeway to compensate for delays and that the wintering period generally buffers the variation in timing of autumn migration, but less so when the wintering period is shorter. As such, migrations to more distant wintering areas are expected to be associated with higher risks of delayed spring return in the breeding area - a *net* carry-over effect [24,34].

**Figure 2:**
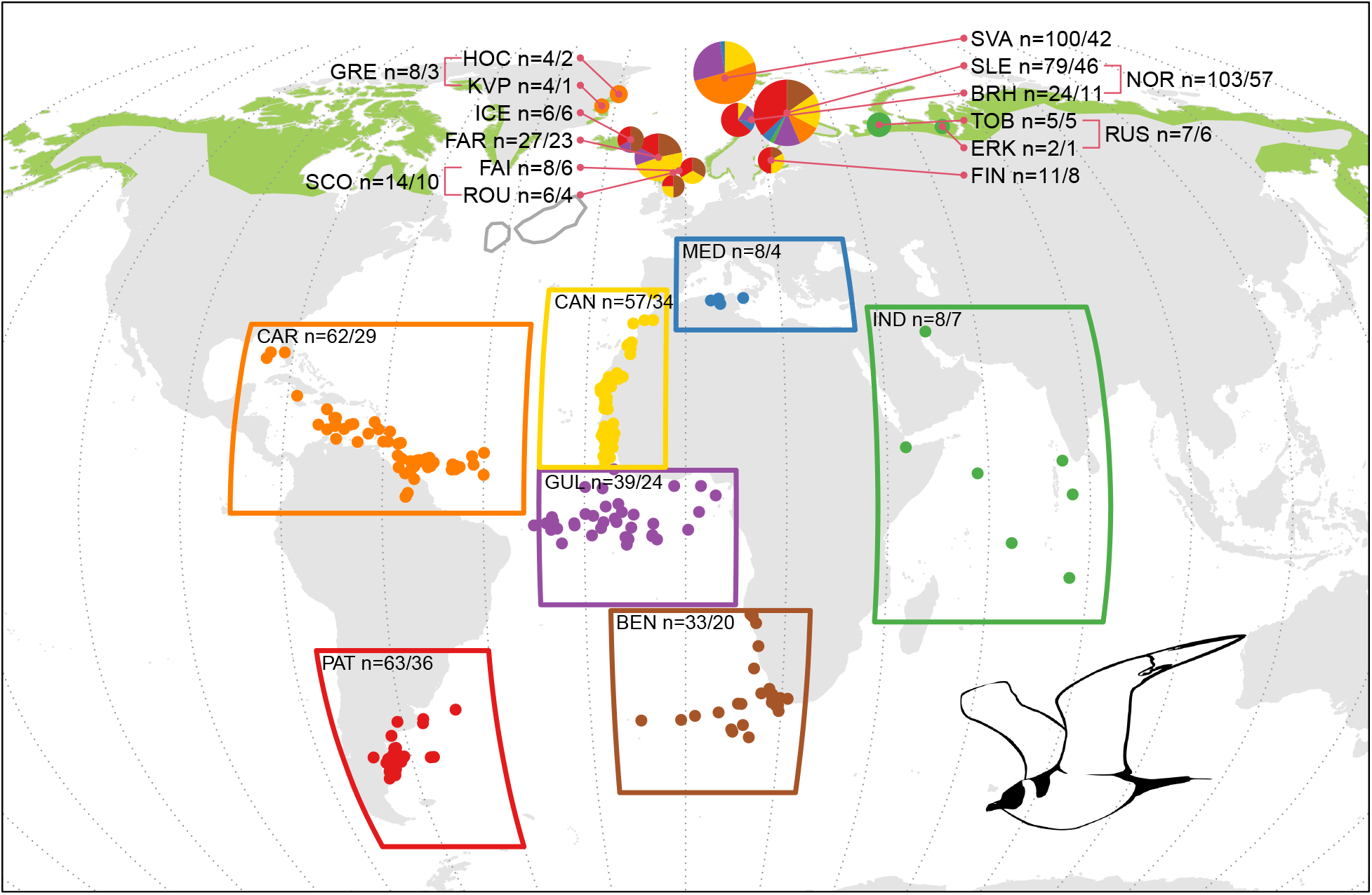
Wintering areas of Arctic Skuas tracked from breeding areas between East Greenland and West Siberia. Dots represent centroids of wintering positions for each track, coloured per wintering area (boxes). Pie charts at breeding sites represent the proportion of individuals wintering in each of the seven wintering areas, with the size corresponding to the total number of individuals tracked. Sample sizes are shown as the number of tracks / number of individuals. Green shaded areas reflect the breeding range. The main stopover in the North Atlantic is outlined in dark grey. Abbreviations for breeding sites are SVA = Svalbard, KVP = Karupelv, HOC = Hochstetter Forland, ICE = Iceland, FAR = Faroe Islands, FAI = Fair Isle, ROU = Rousay, SVA = Svalbard, SLE = Slettnes, BRH = Brensholmen, TOB = Tobseda, ERK = Erkuta and FIN = Finland, and for breeding areas are GRE = Greenland, SCO = Scotland, NOR = mainland Norway and RUS = Russia. Abbreviations for wintering areas are PAT = Patagonian Shelf, BEN = Benguela region, GUL = Gulf of Guinea, CAR = Caribbean region, CAN = Canary Current, MED = Mediterranean Sea and IND = Indian Ocean. Sites within countries were merged for statistical analyses.

## Methods

### Device deployments and sample size

Adult Arctic Skuas were fitted with geolocators (Global Location Sensor or GLS) loggers at twelve breeding sites in eight countries. Data from sites within countries were combined to avoid very small sample sizes in models and are henceforth referred to as breeding areas, as follows: East Greenland (Karupelv, 72°30’N 24°00’W, and Hochstetter Forland, 75°09’N, 19°40’W), Iceland (Hroarstunga, 65°35’N, 14°22’W), Faroe Islands (Fugloy, 62°20’N, 6°19’W), Scotland (Fair Isle, 59°54’N, 1°63’W and Rousay, 59°09’N 3°02’W), mainland Norway (Brensholmen, 69°35’N, 18°01’E and Slettnes, 71°05’N, 28°12’E), Spitsbergen (Kongsfjorden, Svalbard, 79°57’N, 12°06’E), Finland (Satakunta, 61°33’N, 21°27’E), and Russia (Tobseda, 68°36’N, 52°19’E, and Erkuta, 68°14’N, 69°09’E).

Birds were captured either on the nest using bow nets, tent spring traps or walk-in traps, remote release nooses, or away from the nest using net guns. Geolocators were attached to a Darvic ring fitted around the tarsus. Various models of geolocators were used: C65, C250 (Migrate Technology, Cambridge, UK), Mk9, Mk13, Mk15 and Mk18H (British Antarctic Survey, Cambridge, UK) and Mk3006 (Biotrack, Wareham, UK). Geolocators measured ambient light in lux (Migrate Technology) or arbitrary units (BAS/Biotrack) every 1 minute and saved the maximum value every 5 minutes. In subsequent breeding seasons, geolocators were removed and data downloaded.

In total, data for 276 interbreeding periods (i.e. one migration cycle, referred to as a ‘track’ below) from 155 individuals were obtained in 2009-2021. Sample sizes per breeding area ranged from 103 tracks of 57 individuals in mainland Norway to 8 tracks of 3 individuals in East Greenland (figure 2). In total, 34, 23 and 12 individuals were tracked for 2, 3 and 4-6 years, respectively. Some of the tracks (n=48) were incomplete due to battery or electronics failure.

### Geolocator data processing

Positions were estimated for noon and midnight from the light measurements by the geolocators (see electronic supplementary material for details).

We describe the phenology from the perspective of a breeding bird, i.e. ‘autumn migration’ is post-breeding, and ‘spring migration’ is pre-breeding. We define the following key events in the annual cycle: 1) departure from the breeding area; 2) arrival in the wintering area; 3) departure from the wintering area; 4) arrival in the North Atlantic north of 35°N (mostly around the sub-polar frontal zone where many Arctic Skuas made a stopover before returning to the breeding grounds [35,36]); 5) return to the breeding area; 6) clutch initiation. The duration of the intervening periods (in days) were calculated for the: 1) autumn migration; 2) wintering period; 3) spring migration, including the time in the North Atlantic; 4) time in the North Atlantic; 5) pre-laying period; 6) breeding period. As loggers were deployed in the colony and retrieved before departure, a composite index of the length of the breeding period for each track was calculated as the sum of the periods from 1 July (all individuals present in the breeding area) to departure in the first year, and from clutch initiation in the second year to 30 June. Timing of movements were identified by inspecting raw position estimates. For birds breeding in areas with continuous daylight, first position estimates after leaving the zone with continuous daylight were close to the breeding site, but last position estimates, just before returning to the zone with continuous daylight, were around 60°N and distant from the breeding area. To estimate time of departure from or arrival to the breeding area under continuous daylight, we assumed birds traveled at a speed of *ca.* 850 km d^-1^, which has been recorded for the closely related Long-tailed Skua *Stercorarius longicaudus* [37], between the breeding site and the first or last position estimate. Rapid movement to or away from the wintering area, roughly entering or leaving the boxes shown in figure 2, was taken as the timing of arrival at and departure from the wintering area. The date of arrival in the North Atlantic during spring migration was taken as the day birds crossed 35°N. For birds wintering in the Mediterranean Sea, the day after wintering area departure was taken. Clutch initiation dates were inferred from the light data, as incubation stints are apparent as alternating periods of light and darkness each lasting more than an hour [38].

Based on positions in the wintering areas, tracks were assigned to one of seven discrete areas: 1) Caribbean region, Guiyana, Gulf of Mexico and north of Brazil; 2) Patagonian Shelf; 3) Benguela region west to Tristan da Cuñha; 4) Gulf of Guinea and west towards eastern Brazil; 5) Canary Current and off north-west Africa/Iberia; 6) Mediterranean Sea; 7) Indian Ocean and adjacent seas (figure 2). Although skuas moved during the wintering period, sometimes over substantial distances, none had arrival and departure positions in different wintering areas.

### Statistical analysis

Wintering centroids were calculated as the geographical mean of positions between arrival and departure. Minimum migration distance was calculated as the great-circle distance between the wintering centroid and the breeding site. As a measure of migratory connectivity, the degree to which individuals from the same breeding site stay together at wintering areas [30], we calculated the Mantel correlation *R_mantel_* using the ‘MigConnectivity’ package in R [39], using the first track of each individual. Significance of *R_mantel_* was assessed by 1000 random permutations.

Timing of key events and duration of periods were modeled as a function of breeding latitude, with random intercepts for each combination of breeding and wintering area as well as for individuals, in Bayesian Generalized Linear Mixed-effects Models (GLMMs).

We then assessed whether wintering in certain areas led to earlier or later arrival compared to birds from the same breeding area, by constructing GLMMs with breeding and wintering area as fixed effects and random intercepts for individuals. Data with at least three birds for each combination of breeding and wintering areas were selected. We performed two analyses to show the effect of wintering area. First, all pairwise comparisons of wintering area effects within breeding areas were made based on the posterior distributions of the contrast between each set of two wintering areas. Posterior contrast probabilities where 0 was outside the 95% high density interval (HDI) were considered important. Second, to test whether wintering areas led to advances or delays relative to birds from the same breeding area but wintering elsewhere, the 95% HDI of the posterior distributions of each breeding-wintering area combination were compared to the region of practical equivalence (ROPE), defined as the mean timing of birds from the same breeding area but wintering elsewhere, ± 0.1 * sd [40]. If the 95% HDI of the posterior distribution fell completely outside the ROPE, the wintering area was taken to have an important effect. Variance components of the fixed and random effects were used to show partitioning of variation between and within individuals (see supplementary material).

Using only data from breeding areas with the largest data sets (FAR, NOR and SVA), we tested whether the duration of periods between events changed according to timing of the onset of each period, by fitting GLMMs with duration as the response, the onset of the period as a fixed effect, random intercepts and slopes for breeding-wintering area combinations and random intercepts for individuals. To test whether the duration of periods also changed as a function of timing within each breeding-wintering area combination, GLMMs were constructed with duration as the response, slopes with timing per breeding-wintering area as fixed effects and random intercepts for individuals.

GLMMs were fitted using the brms package in R [41], which provides an interface to stan [42], with 6000 iterations, a burn-in of 2000, a thinning interval of five and default uninformative priors. Model results were checked for mixing of and autocorrelation within the chains, and for convergence (*R̂* ∼ 1).

## Results

### Migratory connectivity and spatial consistency of individuals

For any given colony, there was a range of wintering areas used by individual Arctic Skuas (figure 2). The largest variation was among individuals from Slettnes, mainland Norway (largest sample size, n=46 individuals), which wintered in all seven areas identified in this study. Variation among individuals was smallest in the small samples from Russian sites that all wintered in the Indian Ocean (n=6), and from Greenland, that all wintered in the Caribbean region (n=3). Arctic Skuas breeding in Svalbard (n=42) wintered primarily in the Caribbean region (51%), and none migrated to the southern wintering areas, the Patagonian Shelf and Benguela region. In contrast to the Svalbard population, few individuals from mainland Norway wintered in the Caribbean region (9%, n=57). A considerable proportion of individuals breeding in mainland Norway and Finland wintered on the Patagonian Shelf (43%, n=63). Birds breeding in Iceland, the Faroe Islands and Scotland wintered mainly in the Benguela region (31%), Canary Current (36%) and the Patagonian shelf (23%, n = 39), but none in the Caribbean or Mediterranean Sea. Individual wintering-site fidelity was very high (supplementary material).

The considerable degree of mixing of individuals from different breeding sites in wintering areas indicated weak overall migratory connectivity, as is expressed in the *R_mantel_* of 0.19 (95% HDI: 0.11 – 0.26); excluding the birds from the Russian sites resulted in a slightly lower *R_mantel_* of 0.14 (95% HDI: 0.07 – 0.22).

During both autumn and spring migration, most birds wintering in the Atlantic staged in a broad area across the central North Atlantic, roughly between 35°N and 55°N (figure 2). Individuals breeding at Svalbard and wintering in the Canary Current, however, skipped this staging area in spring and followed a more eastern route along Western Europe [36].

### Annual schedules in relation to migration strategies

Birds at more northerly colonies started breeding (i.e. laid the first egg) later (figure 3, supplementary material, figure S1) and also started and finished migration later. The duration of spring migration, the time spent in the North Atlantic and the time between return to the breeding area and clutch initiation all decreased with breeding latitude (figure 3). However, the duration of autumn migration, wintering and breeding periods did not change with breeding latitude (figure 3).

**Figure 3:**
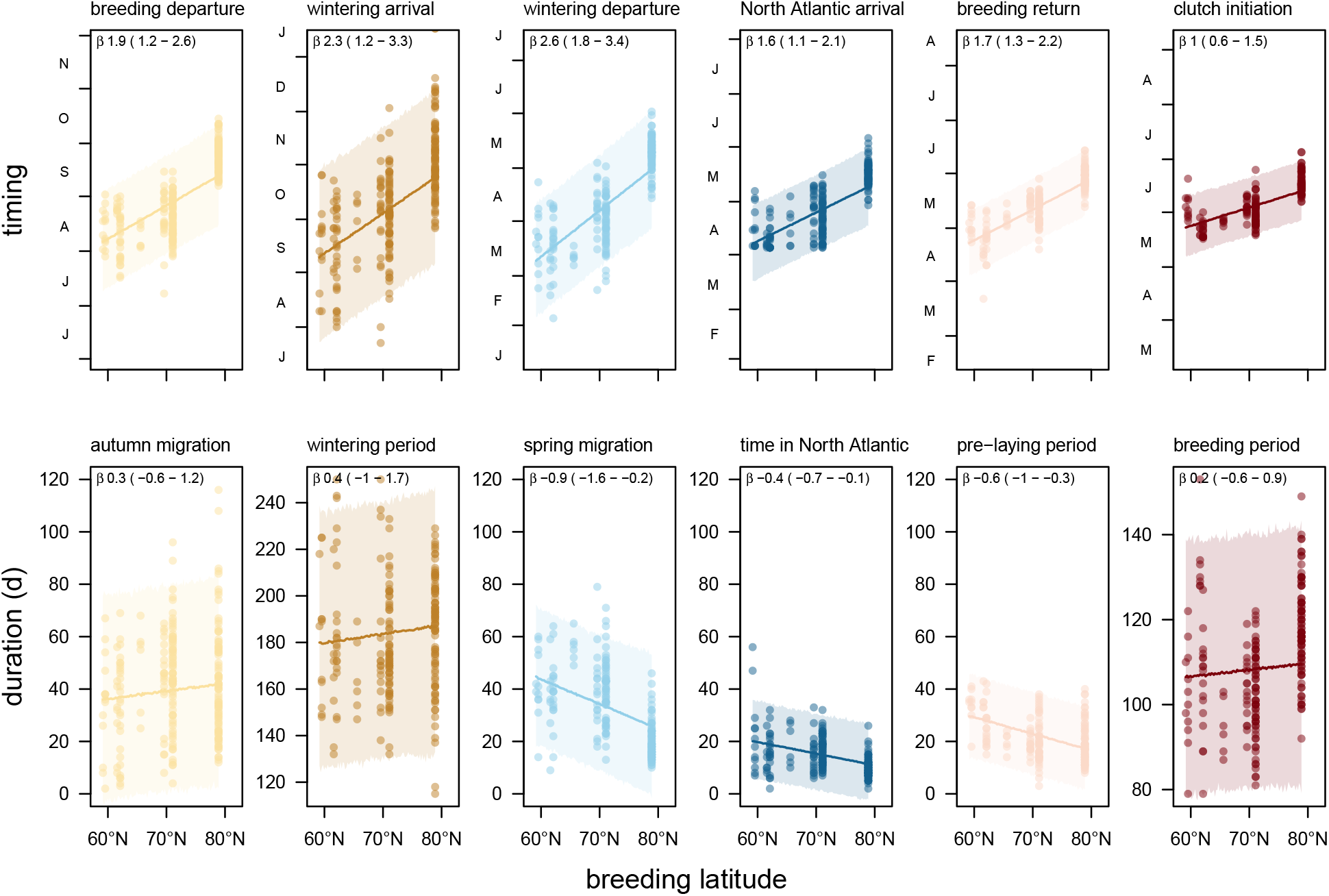
Relation between breeding latitude and timing (upper panels) or duration (lower panels) in Arctic Skuas tracked from breeding areas between East Greenland and West Siberia to wintering areas in the Atlantic. Shaded areas are 95% HDIs.

Birds that used different wintering areas also showed differences in timing and duration of migration (figure 4). There was a clear trend of longer migration periods and shorter wintering period when wintering further south for all breeding areas. Birds wintering on the Patagonian Shelf and the Benguela region (the two furthest wintering areas, at 12647-14664 km and 9803-11800 km, respectively, from colonies) generally stayed for shorter periods in the wintering areas and had longer spring migrations than individuals wintering in the Canary Current (the closest wintering area, at 4021-7140 km from colonies), with the Guinean and Caribbean regions showing intermediate duration of migration and wintering periods. Birds from the same breeding population returned to the North Atlantic at the same time regardless of wintering area, with the exception of later timing among birds wintering on the Patagonian Shelf, and earlier timing among Norwegian birds wintering in the Canary Current. In all breeding areas except Norway, the timing of return to the breeding area was similar between birds from different wintering areas. Among Norwegian birds, we found weak evidence that return to the breeding grounds was slightly delayed among birds that wintered on the Patagonian Shelf and was advanced among those that wintered in the Canary Current. Although timing of clutch initiation varied among Norwegian birds — birds that wintered in the Caribbean region bred substantially earlier — there was no relation between wintering area and clutch initiation date across breeding areas. From wintering area arrival to clutch initiation in the next breeding season, variation between and within individuals declined (electronic supplementary material, figure S2).

**Figure 4:**
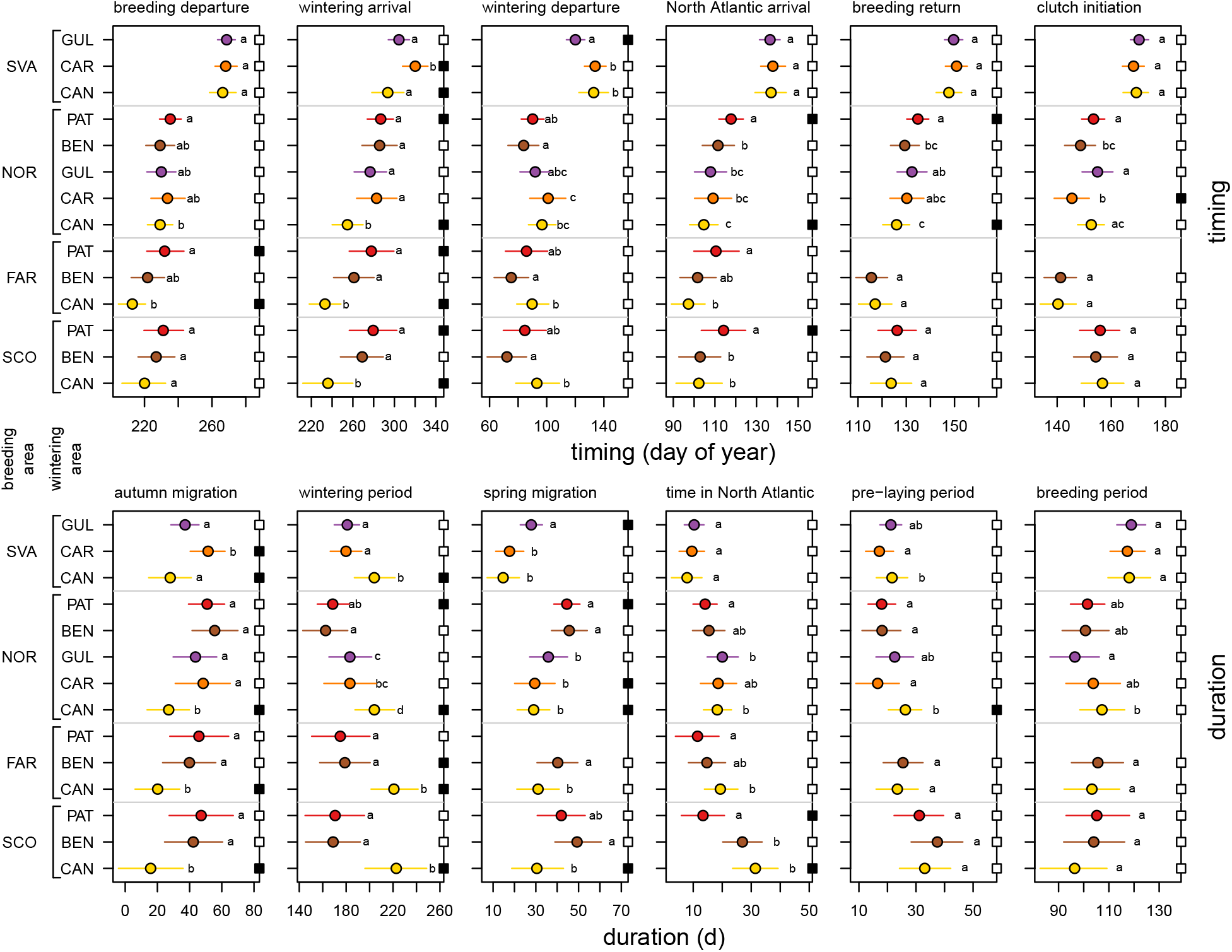
Parameter estimates of the timing of events (upper panels) or duration of phases (lower panels) per breeding and wintering area for Arctic Skuas tracked from breeding areas in the North Atlantic to wintering areas in the Atlantic. Breeding areas are ordered from South (SCO) to North (SVA) and wintering areas from North (CAN) to South (PAT). Error bars show 95% HDIs. Within breeding areas, estimates accompanied by a common letter are equivalent based on whether 0 was included in the 95% HDI of their contrast posterior distribution (values are in supplementary material table S1). Filled squares on the right y-axes indicate that the 95% HDI was outside the ROPE of birds from the same colony wintering elsewhere. Colours correspond to the wintering areas in figure 2.

### Strength of temporal carry-over effects

Both between and within breeding populations, the duration of each migration phase was shorter when started later (figure 5). There was no consistent pattern among breeding areas in the steepness of this relationship, and slope estimates (a measure of the *strength* of the carry-over effect) were not consistently associated with breeding or wintering areas (electronic supplementary material, figure S3). Overall, the decline of duration with time was steepest for the wintering period (electronic supplementary material, figure S3b), when slope estimates around -1 indicate complete compensation of the timing of arrival in the wintering area (figure 1). There were two exceptions to this pattern: slope estimates around -0.5 for the two farthest wintering areas suggested only partial compensation for late arrival in a) Norwegian birds wintering at the Patagonian Shelf (*β* = -0.5, 95% HDI = -0.83 – -0.15) and b) Faroese birds wintering in the Benguela region (*β* = -0.48, 95% HDI = -1.2 – 0.24). Note, however, that the estimate for the Faroese birds wintering in the Benguela region was associated with a very wide HDI overlapping with both -1 and 0, and that the slope estimate for birds from mainland Norway wintering in the Benguela region indicates complete compensation (*β* = -0.97, 95% HDI = -1.53 – -0.43). The duration of spring migration and the time spent in the North Atlantic were tightly related to their timing, with overlapping distributions across breeding and wintering areas (figure 5c-d). The slope estimates between -0.5 and 0 indicated a partial to complete carry-over effect of the arrival in the North Atlantic on return to the breeding area (figure 5d-e). The timing of return partially carried-over to the timing of clutch initiation, with most slope estimates between -1 and -0.5 (electronic supplementary material, figure S3e).

**Figure 5:**
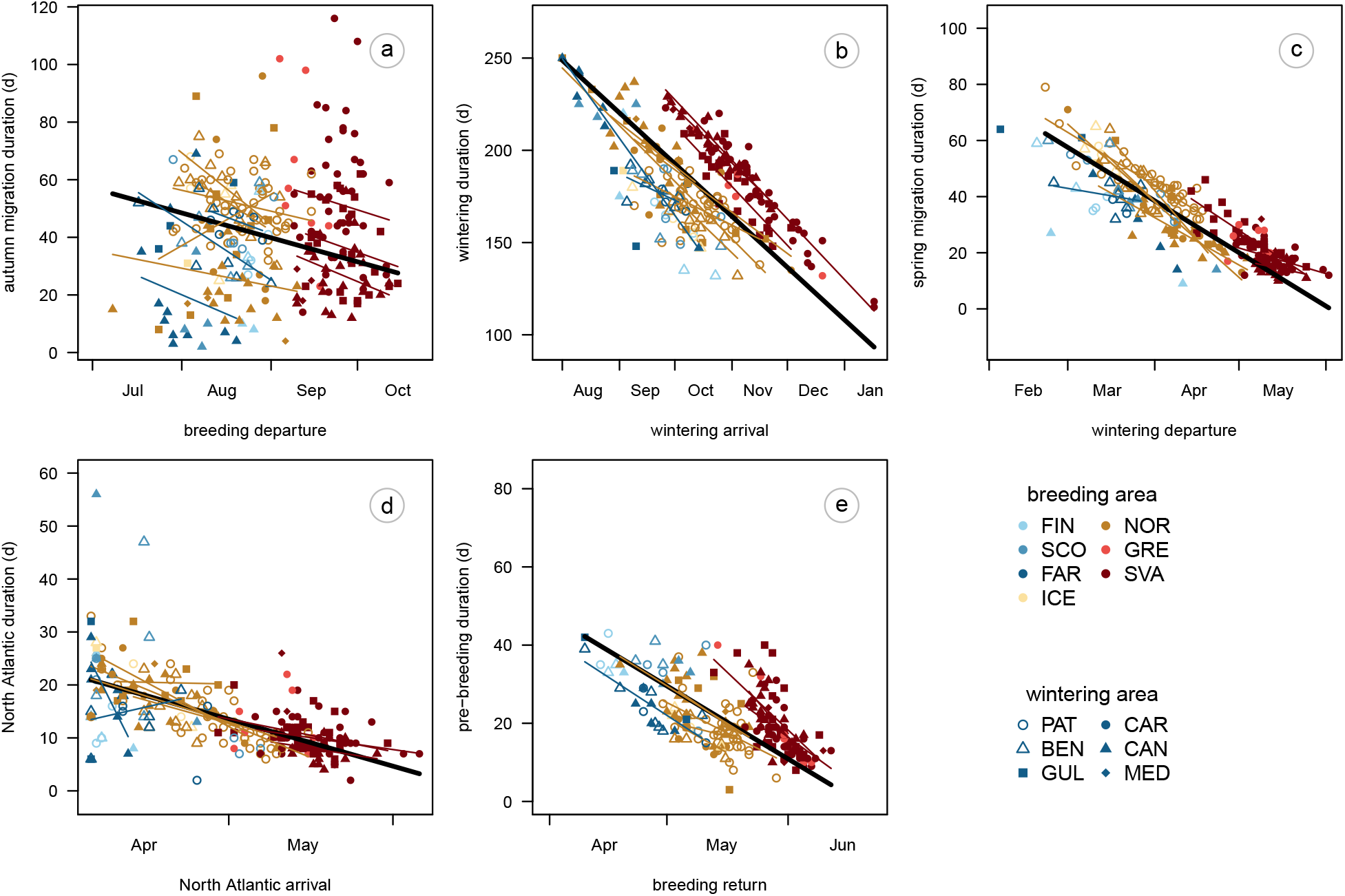
The relationship between duration of each period and the timing of its start in Arctic Skuas tracked from breeding areas in the North Atlantic to wintering areas in the Atlantic. Thick lines show overall slope estimates, thin lines show regression lines per combination of breeding and wintering area. Note that the aspect ratio of all plots is 1, but the axis scales differ.

## Discussion

Our study demonstrates substantial variation within and between breeding populations in both wintering areas and annual schedules of the Arctic Skua, a long-distance migratory seabird with a huge wintering range. Arctic Skuas showed weak overall migratory connectivity in the Atlantic, with very high variation among individuals, including individuals within breeding sites, in their use of wintering areas separated by thousands of kilometers. Individuals from the colony with the most tracked birds (Slettnes, mainland Norway) wintered in all productive areas of the Atlantic between 50°S and 30°N, and also in the Mediterranean Sea and the Persian Gulf (figure 2), together representing roughly a tenth of the world’s ocean surface area. The lower variability in wintering destinations of birds from other breeding sites may partly reflect smaller sample sizes. The wintering range of Arctic Skuas is as large or larger than most other well-studied long-distance migratory seabirds [43–46] and also far exceeds the variation shown by a congeneric species, the Long-tailed Skua *Stercorarius longicaudus* [47]. Such ocean-scale variation in wintering area selection makes the Arctic Skua exceptionally well-suited for directly testing the relationships between the timing of annual events and wintering site selection within and between breeding colonies.

Migration timing of the tracked Arctic Skuas varied principally with breeding latitude, with later-timed schedules of more northern breeders associated with faster spring migrations and shorter periods between spring arrival at the breeding grounds and clutch initiation. Variation in timing of autumn migration was completely compensated during the wintering period, even in the most distant wintering areas, where skuas spent less time. Despite the large variation in migration distances and schedules and the *strong* carry-over effect of timing during spring migration, the timing of return to the breeding area was highly synchronized among individuals from the same site, leaving no substantial *net* carry-over effect.

In Arctic Skuas, breeding latitude not only shifted the timing of annual schedules, as expected [12,15,16,48], but later timing also affected the duration of subsequent key periods within the annual cycle. Most notably, Arctic Skuas breeding at high latitudes had faster spring migrations and shorter pre-laying periods compared to birds breeding at lower latitudes. The faster migration of later-timed spring migration of Arctic Skuas contrasts with little or no geographical variation in the duration of spring migration of two insectivorous songbirds [16,48] and a long-distance migratory seabird, the Arctic Tern *Sterna paradisaea* [49]. Arctic Skuas may face faster seasonal changes in conditions encountered *en route* or may be more flexible in where they fuel for migration/reproduction, by regulating the amount of pre-migratory fueling in the wintering area, foraging *en route* and the time spent in staging areas, in particular the North Atlantic staging area where Arctic Terns stage for no or few days in spring [49]. Arctic Skuas from Svalbard spent fewer days during spring migration at stopovers compared to birds from more southern breeding areas (figure 5d; [36]), which may be explained by deteriorate foraging conditions in the North Atlantic for Arctic Skuas during April to May, when the potential host species to kleptoparasitize [50], such as Arctic Terns and Sabine’s Gull *Xema sabini*, return to their breeding areas [49,51,52]. Shorter time spent at stopovers suggests that birds from more northern breeding areas need to fuel up before migration rather than depend on foraging *en route* to fuel the migration flight, which is common among migrating seabirds [53], and to return with large body reserves in the breeding area [54]. These body stores are also used for egg production: in a population breeding at 61°N, a latitude similar to our most southern study sites and thus presumably with a relatively long pre-laying period, 16% of the protein in eggs originated from distantly-acquired resources [55]. Carrying large nutrient stores on migration is however costly and hence less beneficial as migratory distance increases. This trade-off potentially explains why Arctic Skuas breeding in the high Arctic do not migrate to the southernmost wintering areas. However, carrying large nutrient stores on migration can also be advantageous as it increases flight speed when adopting the ‘dynamic soaring’ flight technique [56], which skuas utilize at high wind speeds. Furthermore, carrying nutrient stores is relatively less costly in larger-bodied birds [19]. Interestingly, Arctic Skuas from Svalbard were heavier and larger than those from mainland Norway, the Faroe Islands and Scotland (B. Moe, R.S.A. van Bemmelen, K.R.S. Snell, R.A. Phillips *unpublished data*). Selection for larger body size to reduce the relative costs of carrying nutrient stores during migration provides an alternative hypothesis to explain latitudinal clines in body size, for which Bergman’s rule is usually invoked [57].

Arctic Skuas were able to compensate for the timing of the autumn migration during the wintering period. Thereby, our study adds to a growing number of studies of ducks, waders and passerines reporting the wintering period to be the main period in the annual cycle where delays are compensated [17,27,58,59], although the migration periods may additionally act as time buffer in some species [60,61]. The ability to shorten the wintering period to compensate for later timing of autumn migration has also been shown by experimentally increasing the duration of parental care [62]. We investigated whether birds are less able to compensate for timing of autumn migration if the wintering period is shorter, as observed in more distant wintering areas, and evidence was inconsistent. Indeed, reduced compensation in the wintering period was found among skuas breeding in northern Norway and wintering at the furthest wintering area, the Patagonian Shelf (figure 4), where the wintering period is up to 50 d shorter than in the northernmost wintering area (but still about 50 d longer than the 122 d needed for the moult of the primary feathers; [63]). However, no conclusive evidence for reduced compensation was found among individuals wintering in the Benguela region, at a similar latitude and migration distance. Therefore, site-specific effects, rather than migration distance alone, may explain the difference in the degree to which birds compensate earlier timing variation during the wintering period, which corresponds to findings for the Sanderling *Calidris alba* [13].

Our study shows largely synchronized timing of return to the breeding grounds as well as clutch initiation within breeding sites, despite differences between wintering areas in migration distance of thousands of kilometers and in average spring migration duration of up to 19 d (in Scotland). Despite the longer duration of spring migration from more distant wintering areas and the increasing *strength* of the carry-over effect of timing during spring migration, the similar departure dates across wintering areas did not result in substantial *net* carry-over effects on the breeding grounds. This indicates that, rather than departing earlier from the wintering grounds, Arctic Skuas with longer migrations increased travel speeds to ensure a timely return to the breeding grounds, which agrees with empirical data from songbirds [21]. The similarity in timing of departure across wintering areas observed in Arctic Skuas suggests control by endogenous biological ‘clocks’, which are thought to be fine-tuned by selection [31].

Birds from widely separated wintering areas arrived back in the breeding area, on average, within less than a week. This synchrony underpins the strong selection on timing of spring arrival and breeding when competition for territories is high [64]. Later arrival at the breeding grounds of individuals from more distant wintering areas has been reported for several short- to long-distance migrants [28,65–69], but only in European Shags *Gulosus aristotelis* and Eurasian Spoonbills *Platalea leucorodia* did this also lead to later reproduction (6 d in shags, 12 d in male spoonbills) and lower reproductive success. In Arctic Skuas, however, we found partial compensation for later return at the breeding grounds: on average, a 1 d later spring arrival led to 0.6 d later clutch initiation, but with substantial individual variation. We did not have data on reproductive success, but as wintering area had a minimal influence on timing of return to the breeding area (on average up to 6 d later) and laying dates (up to 3 d), a substantial effect of wintering area on reproductive success via timing of breeding seems unlikely. Furthermore, unless an effect on productivity is substantial, demonstrating a carry-over effect on reproduction would be difficult considering the multitude of local effects at the breeding grounds – effects that have been shown to impact reproduction and thus drive population declines in Arctic Skuas, at least in Scotland and mainland Norway [70,71].

By describing the migratory connectivity and ocean-scale geographic variation in annual schedules and carry-over effects of a long-distance migratory seabird, our study contributed to our understanding of potential drivers of meta-population dynamics in migratory species. Our results indicate that choice of wintering area is unlikely to affect population trends through a carry-over effect on timing of breeding, irrespective of breeding latitude. Of course, this does not rule out effects on survival or reproduction via other mechanisms. For example, as daily survival rate may differ between stages of the annual cycle [7,8], wintering area selection may affect annual survival rates and therefore population trends, given their concomitant annual schedules. While the weak migratory connectivity means that conditions in each wintering area can affect multiple breeding populations, each breeding population may be buffered from the effect of conditions at specific wintering areas due the large spread of individuals across wintering areas [30]. Interestingly, the population decline in Svalbard (from where no individuals wintered in the two most distant wintering areas) is less than in the Faroe Islands, Scotland and mainland Norway [70–72] and the longevity record holder of the species, a bird from Finland, wintered in one of the closer wintering areas, the Canary Current [73]. If and how breeding area, wintering area, and the corresponding annual schedules affect seasonal survival rates in Arctic Skuas, and whether these can explain differences in population trends, remains to be investigated.

## Declarations

### Ethics

This study was carried out in accordance with guidelines in the use of wild birds for research [74]. Catching and tagging Arctic Skuas was approved for the following countries by the appropriate licensing authority: East Greenland (Government of Greenland, #C11-31, C12-28, C13-4(29), C13-11, C14-4(23), C14-16, C15-4(10), C15-14, C16-4(15), C16-20, C17-3(28), C17-23), Svalbard/Norway (Governor of Svalbard and Norwegian Food Safety Authority, FOTS ID 2086, 3817, 6329, 8538, 15726); Iceland (Icelandic Bird Ringing Scheme); Faroe Islands (Copenhagen Bird Ringing Centre and The National Museum of the Faroe Islands); Scotland (Special Methods Technical Panel of the British Trust for Ornithology Ringing Committee); Finland (Centre for Economic Development, Transport and the Environment, VARELY/418/07.01/2012) and Russia (Federal Law No. 52-FZ of April 24, 1995, Article 44).

### Data accessibility

Geolocator data is stored on www.movebank.org under study IDs: 2746281685 (Svalbard, Norway), 968984187 (Slettnes, Norway), 2746341928 (Brensholmen, Norway), 976692287 (Tobseda, Russia), 1260558178 (Greenland), 2746320914 (Iceland), 2746186408 (Faroe Islands), 2743674038 (Scotland) and 2747912295 (Finland). The studies can also be found by searching for the study IDs using the search functionality at this website.

### Consent for publication

Not applicable.

### Author contributions

conceptualization: R.S.A.vB., B.M., H.S., K.R.S.S. & I.T.; methodology, formal analysis & visualization: R.S.A.vB.; fieldwork: R.S.A.vB.; B.M., H.S., S.A.H. K.R.S.S., S.H., E.M., K.N., M.S., D.E., S.R.V., G.T.H., Y.K., S.H., J.C., O.G., J.L., B.S., A.S., I.T. writing original draft: R.S.A.vB. with input from B.M., H.S., I.T., K.R.S.S., E.M.H & R.P.; writing and editing: R.S.A.vB., B.M., H.S., I.T., K.R.S.S., E.M.H, J.C. & R.P.; supervision: B.M. & I.T.; funding acquisition: R.S.A.vB., B.M., E.M.H, J.C., E.M. & I.T.

## Acknowledgements

This study would not have been possible without the help of the many fieldworkers and others that invested substantial time and resources. In Slettnes, we were assisted in the field by Geert Aarts, Daniël van Denderen, Jan van Dijk, Maria van Leeuwe, Daan Liefhebber, Morrison Pot, Marc van Roomen, Janne Schekkerman and Rinse van der Vliet. Other help was provided by Cees and Mimi Tesselaar, Barbara Ganther, Hans-Ulrich Rösner, Karl-Birger Strann, Jeanette Hickman, Torstein Johnsrud, Niels Westpahl and Benedicte Færevåg. In Svalbard and Brensholmen, help was provided by Erlend Lorentzen, Elise Biersma, Fokje Schaafsma, Anouk Goedknegt, Oebele Dijk, Maarten Loonen, Anette Fenstad, Elise Skottene, Liv Monica Trondrud, Nora Bjørnlid, Heidi Kilen, Eline Rypdal, Emilly Hill, Melissa Fontenille and Thomas Oudman. Expeditions to Tobseda in 2014-2015 were organized by Thomas Lameris and in 2018 by Götz Eichorn (NIOO-KNAW), and joined by Kees Schreven, Stefan Sand, Jasper Koster and Chiel Boom. In the Faroe Islands, help was provided by Jón Aldará, Leivur Janus Hansen and Anthony Weatherhill; we kindly acknowledge the landowners on Fugloy for permission to work on their property. In Greenland, Adrian Aebischer, Vadim Heuacker, Eric Buchel, Brigitte Sabard and Vladimir Gilg helped at Hochstetter, and Anita Lang, Mark Nitze and Felix Norman at Karupelv Valley. In Erkuta, help was provided by Natalia Sokolova, Ivan Fufachev and Stijn Hofhuis. In Finland, Kari Mäntylä and Jukka Nuotio assisted during fieldwork. Support in the field in Scotland was provided by Helen and David Aiton on Rousay and the Fair Isle Bird Observatory. James Fox (Migrate Technology Ltd) and Glen Fowler (Biotrack Ltd) kindly retrieved data from loggers that failed to download.

