## supplementary material for "Ocean-scale variation in migration schedules of a long-distance migratory seabird is fully compensated upon return to the breeding site"

### Posterior probabilities

Table S1. Posterior probability that the difference in timing or duration between two wintering areas when originating from the same breeding area is more extreme than 0 in Arctic Skuas tracked from breeding areas in the North Atlantic to wintering areas in the Atlantic. Posterior probabilities of 0.1 and lower are highlighted in red, with more intense red for lower values.

| breeding area | wintering areacomparsion | breeding departure | wintering arrival | wintering departure | North Atlantic arrival | breeding return | clutch initiation | autumn migration | wintering period | spring migration | time in North Atlantic | pre-laying period | breeding period |
|---|---|---|---|---|---|---|---|---|---|---|---|---|---|
| SVA | CAR vs. GUL | 0.43 | 0.01 | 0.00 | 0.33 | 0.31 | 0.16 | 0.01 | 0.44 | 0.00 | 0.37 | 0.06 | 0.33 |
| | CAN vs. GUL | 0.27 | 0.09 | 0.01 | 0.44 | 0.24 | 0.34 | 0.09 | 0.01 | 0.00 | 0.18 | 0.44 | 0.43 |
| | CAN vs. CAR | 0.31 | 0.00 | 0.42 | 0.41 | 0.10 | 0.30 | 0.00 | 0.00 | 0.20 | 0.24 | 0.03 | 0.42 |
| NOR | BEN vs. PAT | 0.06 | 0.44 | 0.10 | 0.04 | 0.02 | 0.04 | 0.23 | 0.24 | 0.38 | 0.30 | 0.48 | 0.43 |
| | GUL vs. PAT | 0.09 | 0.08 | 0.36 | 0.00 | 0.20 | 0.29 | 0.12 | 0.05 | 0.02 | 0.01 | 0.07 | 0.14 |
| | CAR vs. PAT | 0.36 | 0.34 | 0.03 | 0.02 | 0.08 | 0.00 | 0.38 | 0.08 | 0.00 | 0.06 | 0.34 | 0.31 |
| | CAN vs. PAT | 0.04 | 0.00 | 0.07 | 0.00 | 0.00 | 0.35 | 0.00 | 0.00 | 0.00 | 0.03 | 0.00 | 0.07 |
| | GUL vs. BEN | 0.44 | 0.16 | 0.09 | 0.20 | 0.20 | 0.04 | 0.07 | 0.03 | 0.02 | 0.08 | 0.13 | 0.23 |
| | CAR vs. BEN | 0.21 | 0.40 | 0.01 | 0.31 | 0.42 | 0.19 | 0.22 | 0.04 | 0.00 | 0.19 | 0.36 | 0.29 |
| | CAN vs. BEN | 0.50 | 0.00 | 0.01 | 0.04 | 0.14 | 0.10 | 0.00 | 0.00 | 0.00 | 0.15 | 0.01 | 0.09 |
| | CAR vs. GUL | 0.25 | 0.27 | 0.09 | 0.40 | 0.29 | 0.01 | 0.30 | 0.50 | 0.12 | 0.34 | 0.08 | 0.11 |
| | CAN vs. GUL | 0.44 | 0.00 | 0.20 | 0.18 | 0.03 | 0.23 | 0.01 | 0.02 | 0.07 | 0.23 | 0.14 | 0.02 |
| | CAN vs. CAR | 0.19 | 0.00 | 0.25 | 0.17 | 0.11 | 0.01 | 0.01 | 0.04 | 0.46 | 0.45 | 0.01 | 0.26 |
| FAR | BEN vs. PAT | 0.06 | 0.10 | 0.11 | 0.08 | | | 0.29 | 0.39 | | 0.23 | | |
| | CAN vs. PAT | 0.00 | 0.00 | 0.32 | 0.01 | | | 0.01 | 0.00 | | 0.03 | | |
| | CAN vs. BEN | 0.05 | 0.00 | 0.02 | 0.19 | 0.36 | 0.40 | 0.01 | 0.00 | 0.05 | 0.10 | 0.34 | 0.37 |
| SCO | BEN vs. PAT | 0.28 | 0.24 | 0.08 | 0.04 | 0.16 | 0.38 | 0.34 | 0.44 | 0.14 | 0.00 | 0.13 | 0.44 |
| | CAN vs. PAT | 0.08 | 0.00 | 0.20 | 0.04 | 0.31 | 0.43 | 0.01 | 0.00 | 0.06 | 0.00 | 0.37 | 0.14 |
| | CAN vs. BEN | 0.19 | 0.01 | 0.01 | 0.46 | 0.32 | 0.32 | 0.02 | 0.00 | 0.00 | 0.17 | 0.22 | 0.16 |

### Individual consistency

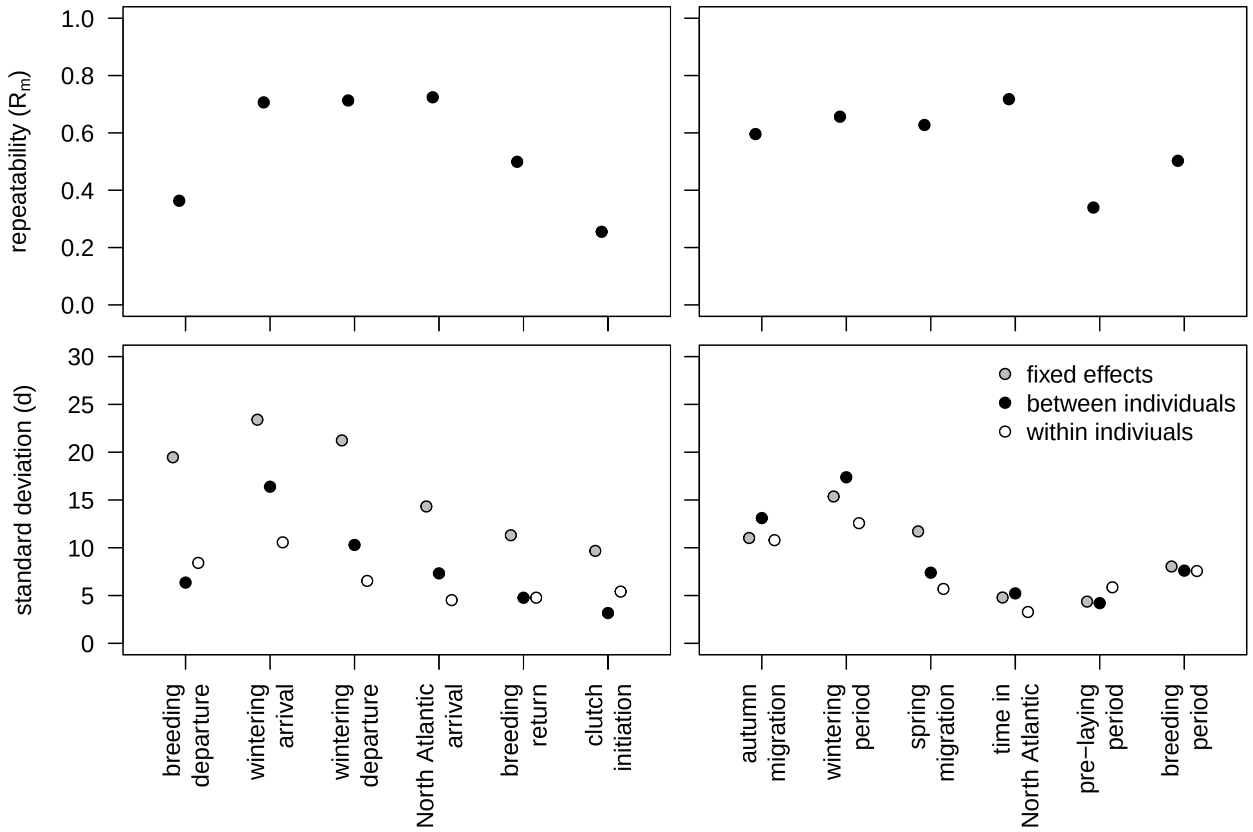


Figure S1. Marginal repeatability (upper panels) and squared posterior means of model variance components of GLMMs of timing and duration (lower panels) of timing of events (left) and duration of phases (right) in the annual cycle of Arctic Skuas tracked from breeding areas in the North Atlantic to wintering areas in the Atlantic.
